# The Smac mimetic drug Xevinapant sensitizes triple negative breast cancer to radiotherapy-induced cell death in a TNFα dependent manner

**DOI:** 10.64898/2026.09.29.755558

**Authors:** Mame Daro Faye, Julie Lafontaine, Philip Wong, Cynthia Ménard

**Affiliations:** Department of Radiation Oncology, Centre Hospitalier de l’Université de Montréal, Montréal, Canada; Centre de Recherche du Centre Hospitalier de l’Université de Montréal (CRCHUM), Montréal Canada; Institut du Cancer de Montréal, Montréal, Canada; Department of Radiology, Radiation Oncology and Nuclear Medicine, Faculty of Medicine, Université de Montréal, Montréal, Canada; Radiation Medicine Program, Princess Margaret Hospital and University of Toronto, Toronto, Canada

**Keywords:** Radiotherapy, Xevinapant, Smac mimetics, TNFα, TNBC, Radiosensitization

## Abstract

**Purpose:** There is an unmet need in the treatment of triple-negative breast cancer (TNBC), as recurrence rates remain high; thus, novel treatment strategies are warranted to improve outcomes for patients. Smac mimetics (SMs) are small molecules that inhibit the inhibitors of apoptosis. In this study, we investigated the properties of the SM Xevinapant (AT406) as a radiosensitizer in TNBC.

**Methods:** TNBC cell lines were treated with Xevinapant (AT406) in the presence or absence of ionizing RT and cell viability and apoptosis measured *in vitro*. Clonogenic assays were performed to confirm radiosensitization. We also assessed xevinapant radiosensitization in the presence of the Tumor Necrosis Factor α (TNFα) cytokine, which has been shown to enhance SM-induced apoptosis. Radiosensitization mechanisms were further validated by western blotting, flow cytometry and cell cycle analysis.

**Results:** We show that Xevinapant increased cell death in a panel of irradiated TNBC cell lines. In clonogenic assays, Xevinapant sensitized MDA-MB-231 cells to RT-induced cell death, and this was significantly enhanced in the presence of TNFα, with a dose-enhancing factor (DEF) of 1.57. We also show that Xevinapant radiosensitization is mediated by TNFα-dependent apoptosis.

**Conclusions:** Xevinapant is a radiosensitizer that enhances RT-induced cell death in TNBC cells, and TNFα is central to this process through activation of apoptosis. These results are promising for combining RT with Xevinapant to improve local control in TNBC.

## Background

Triple negative breast cancer (TNBC) is an aggressive type of breast cancer with poor prognosis. Despite advances in therapeutic options, the locoregional and distant failure rates in TNBC remains high and are a major cause of death.^1^ Indeed, treatment of TNBC represents a challenge due to the paucity of targetable pathways as opposed to estrogen receptor-positive or Her2-positive breast cancers. In recent years, the use of PARP inhibitors and of immunotherapy have shown promising results in TNBC,^2,3^ however there is room for improvement as recurrence rates can be as high as 50% in patients that did not achieve a pathological complete response (pCR).^4^ Thus, novel treatment strategies are warranted to improve outcomes for patients with TNBC.

Radiotherapy (RT) remains one of the mainstays of adjuvant therapies for breast cancer, decreasing disease recurrence by 50% and cancer-specific mortality by 20%.^5^ RT works by creating double-stranded breaks in DNA leading to mitotic catastrophe and cell death.^6^ Apoptosis is one of the mechanisms by which damaged irradiated cells are eliminated and one of the hallmarks of cancer cells is their ability to escape apoptosis, thus promoting resistance to various therapies.^7^ The inhibitors of apoptosis (IAPs) cIAP1, cIAP2 and XIAP are major regulators of apoptosis and are overexpressed in many cancers, including TNBC, leading to resistance and poor prognosis.^8^ In particular, cIAP1 and cIAP2 are key inhibitors of Tumor Necrosis Factor α (TNFα)-mediated apoptosis. Indeed, TNFα is a cytokine that binds to its receptor to induce a signaling cascade that will lead to depletion of cIAP1/2 and activation of caspases resulting in apoptosis.^8,9^

SMs are small compound molecules that mimic the second mitochondrial-derived activator of caspases (Smac), which is released from the mitochondria and binds IAPs at the onset of apoptosis, thus allowing it to take place without inhibition.^10,11^ Many groups, including ours, have shown that direct targeting of IAPs by a class of drugs called Smac-mimetics (SMs) induces TNFα-mediated apoptosis.^12-14^ SMs have also been shown to significantly enhance RT-mediated apoptosis and suppress clonogenic survival in a variety of cancers.^15-17^ In recent years, there has been renewed interest in SMs as more clinical trials showed their efficacy in solid cancers in the clinical setting, particularly in head and neck cancer. Indeed, in a randomized phase 2 trial investigating the SM Xevinapant plus chemoradiotherapy (CRT) vs. placebo plus CRT in 96 patients with unresected locally advanced head and neck cancer squamous cell cancer (HNSCC), the risk of death was decreased by half in the Xevinapant arm compared with placebo (adjusted HR 0.47; 95% CI, 0.27–0.84; P = .0101) while incidence of late-onset grade ≥3 toxicities was similar across arms^18^.

While preclinical studies suggest the efficacy of smac mimetics in breast cancer, there is a paucity of data in the preclinical and clinical setting to support the use of SMs in combination with radiotherapy as potential therapeutics in breast cancer and particularly TNBC. Combining SMs with RT offers the potential for synergy in inducing apoptosis and increased immunogenic cell death in TNBC. Indeed, SMs have been shown to increase the autocrine production of TNFα, which can trigger the extrinsic apoptotic pathway, complementing the mechanism of action of RT.^6,12^ In this study, we aimed to characterize the potential of the SM drug Xevinapant to potentiate the efficacy of radiotherapy in TNBC. We show that Xevinapant, synergize with RT in the presence of TNFα to induce the death of TNBC cancer cells and that TNFα is central to this process through activation of the programmed cell death.

## Materials and Methods

### Cell culture and expression constructs

Triple negative breast cancer cell lines (MDA-MB-23, MDA-MB-468, BT-549, Hs578T) were purchased from the American Type Culture Collection (ATCC, Manassas, VA, USA). Cells were grown following ATCC recommendations (DMEM or RPMI1640 supplemented with 10% or 20% fetal bovine serum (FBS), with 1% penicillin-streptomycin (Pen-strep). Cells were incubated at 37 °C with 5% CO2 according to ATCC recommendations. Cells were used at a low passage number (lower than 25) upon reception from ATCC, and specifically for less than seven passages for clonogenic assays, in order to maintain their parental phenotype and genotype. Genomic characterization of all investigated cell lines is publicly available.^19^ The NLS-GFP lentiviral construct is a gracious gift from Francis Rodier’s lab. Stable TNBC cell lines expressing the NLS-GFP reporter were established through lentivirus infection, selected through Puromycin selection and maintained in cultured in 0.1% Puromycin containing DMEM or RPMI medium as described above. The NLS-GFP stable cell lines were used in live cell imaging experiments to accurately count the number of living cells (see below).

### Reagents

Xevinapant (AT-406 or Debio 1143) was purchased from Selleckchem (Catalog No S2754), resuspended in dimethyl sulfoxide (DMSO) at a concentration of 1 mmol/L (1 mM) and stored at -20 °C. Final concentrations of Xevinapant were between 1 nM and 1 uM in media, and a final concentration of < 0.1% of DMSO was used in all experiments. Recombinant cell culture grade human tumor necrosis factor alpha (TNFα) was purchased from Enzo Biosciences (Catalog No ALX-520-002), reconstituted in phosphate buffered saline (PBS) and used at a final concentration of 10 ng/ml in media. The pan caspase zVAD-fmk was purchased from MedChemExpress, resuspended in DMSO at a concentration of 20 mM and used at a final concentration of 20 uM in media. For live cell imaging Incucyte experiments, propidium Iodide dye was purchased from Sigma Aldrich (Catalog No P4864-10ML) and the CellEvent™ Caspase-3/7 Detection dye was purchased from Invitrogen (Catalog No C10423).

### Radiation therapy

RT was performed using a Caesium-137 source irradiator (Gammacell 3000 Elan, Best Theratronics, Ottawa, ON, Canada) according to institutional radioprotection guidelines. Cells were seeded in 96-well or 6-well plates and pretreated with drugs as described below before being treated with RT at the indicated doses (between 2 and 8 Gy).

### Live cell imaging

Live cells experiments were performed using the IncuCyte Zoom Live-cell Imaging System (Sartorius, Göttingen, Germany), using a 10X objective. A total of 5 × 10^3^ cells were plated in 96-well plates with 100 μL of medium per well then treated less than 24 hours later with indicated doses of Xevinapant or control DMSO (Time 0). Propidium iodide (PI) staining was used for cell death visualization and was directly added to the media at a final concentration of 1 μg/ml). For Caspase activity experiments, the CellEvent™ Caspase-3/7 reagent was directly added to the media at Time 0 at final concentration of 1:1000. Two or four hours later, cells were irradiated with 8 Gy of ionizing RT vs control (cells removed from incubator but no RT) and then returned to the Incucyte machine for further treatment or monitoring. Two images were taken per well every four hours and cells monitored over 72 to 96 hours. For combination treatments, NLS-GFP stable cells expressing fluorescent GFP were used to be able to quantify more accurately the total number of cells per plate. NLS-GFP/MDA-MB-231 or NLS-GFP/Hs578T cells were plated as described above and treated with 100 nM Xevinapant or DMSO control, TNFα 10 ng/ml or zVAD-fmk 20 μM alone or in combination as indicated. PI or Caspase-37 fluorescent dye were added directly to the media at time 0. Four hours later, cells were treated with 8 Gy of RT and monitored with the Incucyte as described above. Staurosporine 1 μg/ml was used as a positive control for cell death induction.

### Clonogenic survival assays

Cells were seeded in six-well plates at optimized cell densities (100 cells for 0Gy, 100 cells for 2Gy, 200 cells for 4Gy, 800 cells for 6Gy) in 1.5 ml of media and allowed to adhere overnight for 16 hours. Adherent cells were then treated with 1uM of Xevinapant or DSMO 0.1% in a final volume of 2 ml. Four hours later, treated cells were exposed to 0, 2, 4, and 6 Gy of RT using the Gammacell 3000 Elan irradiator. Following treatment, cells were maintained under standard culture conditions for 10–14 days, or until colonies were sufficiently developed. Colonies were then fixed, stained with crystal violet, and colonies containing ≥50 cells were counted. In another set of clonogenic assays, cells were plated and treated similarly but the media changed every 3 days with fresh media containing 1 uM of Xevinapant or control DMSO (q3 days condition). In a third set of experiments cells were irradiated first and two hours later treated with Xevinapant 1 μM or DMSO for 24 hours before the media was changed (Xevinapant followed by RT condition). Plating efficiency (PE) was calculated as *PE = number of colonies formed / number of cells seeded*. To specifically assess for radiosensitization while accounting for any treatment-related effects on baseline clonogenicity, the surviving fraction (SF) at each radiation dose was calculated and normalized to the plating efficiency of the corresponding unirradiated treatment condition as follows: *SF = number of colonies formed / (number of cells seeded × treatment-specific PE)*. Survival curves were obtained by plotting the surviving fractions (log10) as a function of radiation dose. For TNFα-based clonogenic assays, cells were treated with 100 nM Xevinapant, DMSO control or TNFα 10 ng/ml alone or in combination for four hours before exposure to different doses of RT (0, 2, 4, 6 Gy). Caspases neutralization experiments were performed using 20 uM of zVAD.fmk and colony formation assessed as described above.

### Western blotting

Cells were lysed on ice for 30 minutes using a RIPA lysis buffer (Thermo Scientific, Cat No 89900) with 1X protease and phosphatase inhibitors. Total protein content was quantified using a Pierce™ BCA protein assay kit (Thermo Scientific, Cat N0 23225) as directed. Proteins were separated using 4-20% Mini-Protean® TGX Stain-Free precast gels (Biorad, Catalog No 17000435) along with a 10-250kDa Precision Plus protein ladder (Biorad, Catalog No 1610374) and protein transfer performed using the Transblot Turbo nitrocellulose transfer system pack (Biorad, Catalog No 1704159). Antibody staining was performed as described previously^20^ using the following primary antibodies overnight at 4^°^C in 2% bovine serum albumin: rabbit anti-cIAP1 at 1:1000 dilution (Abcam, Catalog No ab108361), mouse anti-cIAP2 at 1:1000 (R&D Systems, Catalog No MAB817), rabbit anti-XIAP at 1:1000 (Abcam, Catalog No ab2541), rabbit anti phospho-cleaved poly ADP-ribose polymerase (phospho-cleaved PARP (Asp214) at 1:1000 (Cell Signalling, Catalog No 12061), rabbit anti-cleaved caspase 3 (Asp175) at 1:1000 (Cell Signaling, Catalog No 9661), rabbit anti-phospho γH2Ax (Ser139) at 1:2000 dilution (Sigma, Catalog No. clone JBW301), anti tubulin (Cell Signaling, Catalog No 2114); and following secondary antibodies: donkey-anti-rabbit IR800 (LI-COR, Catalog No 926-32211) and donkey-anti-mouse IR680 (LI-COR, Catalog No 926-68070) at 1:20 000 dilution. Protein bands were visualized using the Odyssey imaging system and software (LI-COR, Lincoln, Nebraska USA).

### Flow cytometry analysis of apoptosis

Apoptosis was assessed by flow cytometric detection of intracellular active caspase-3. Cells were treated with Xevinapant 100 nM, DMSO control, TNFα 10 ng/ml, zVAD-fmk or combination thereof for four hours before being treated with RT at 4 Gy and incubated for 24, 48 or 72 hours. Adherent cells were collected and washed with phosphate-buffered saline (PBS). In another set of experiments, both adherent and non-adherent cells were collected to recover the total number of cells that underwent apoptosis. Cells were then fixed and permeabilized using the BD Cytofix/Cytoperm™ Fixation/Permeabilization Kit (BD Biosciences, Catalog No 554722) according to the manufacturer’s instructions. Briefly, cells were resuspended in BD Cytofix/Cytoperm™ solution, incubated for 20 min at 4°C, washed with BD Perm/Wash™ buffer, and incubated with BV650-conjugated rabbit anti-active caspase-3 antibody (BD Horizon, Catalog No 5701790) for 30 min at 4°C protected from light. Cells were then washed with Perm/Wash buffer and resuspended in staining buffer for flow cytometric acquisition. Appropriate unstained and fluorescence controls were included to establish gating parameters. Stained cells were counted using the LSR Fortessa™ flow cytometer (BD Biosciences) and analyzed using FlowJo/software v11 (FlowJo, LLC). After exclusion of debris and cell doublets, the percentage of active caspase-3-positive cells was quantified and used as a measure of treatment-induced apoptosis.

### Cell cycle analysis

Cell cycle phase distribution was analysed by flow cytometry. The drug treatments were as described for the combination treatments and four hours later, cells treated with 4 Gy of RT and harvested at various time points. Cells were fixed with 70% ethanol and stained with propidium iodide (Sigma-Aldrich). DNA content for cell-cycle analysis was determined with a LSR Fortessa™ flow cytometer (BD Biosciences).

### Statistical analysis

GraphPad Prism software version 11.0.0 for Windows (Boston, Massachusetts USA) was used for all analyses. Data represents mean +/-standard error of the mean of at least three replicate experiments. Clonogenic survivals, flow cytometry and immunostaining experiments were compared according to the Student’s t-test. Live cell imaging cytotoxicity and apoptosis experiments were compared by two-way ANOVA with Tuckey’s multiple comparison tests. Statistical significance was defined as p<0.05 (95% confidence interval).

## Results

### Xevinapant enhances the radiation sensitivity of TNBC cell lines

A panel of four TNBC cell lines (MDA-MB-231, MDA-MB-468, BT549 and Hs578T) were treated with RT in the presence of increasing concentrations of Xevinapant or control DMSO to assess for differential intrinsic sensitivity to Xevinapant. The different cell line characteristics are shown in Figure 1A. Cells were treated with Xevinapant or control DSMO for four hours followed by 8Gy of RT and cell viability monitored over 72 hours by measuring propidium iodide incorporation into cells. Xevinapant sensitized three of the four cell lines to RT at varying concentrations, and the MDA-MB-231 cell line showed the most sensitivity. Indeed, at 72 hours, Xevinapant increased RT-induced cytotoxicity in MDA-MB-231 by 14-fold at 100 nM and by 30-fold at 1uM compared to control (DMSO at T=0) (Figure 1B), by 16- and 17-fold respectively in MDA-MB-468 (Figure 1C), by 10- and 13-fold respectively in BT549 (Figure 1D) and by 5- and 7-fold in Hs578T (Figure 1E). The MDA-MB-468 cell line (Figure 1B) and Hs578T cell line (Figure 1E) exhibited substantial cell death over 72 hours even under DMSO control conditions, likely reflecting inherent sensitivity to the growth conditions and/or to increased confluence over time.

**Figure 1.**
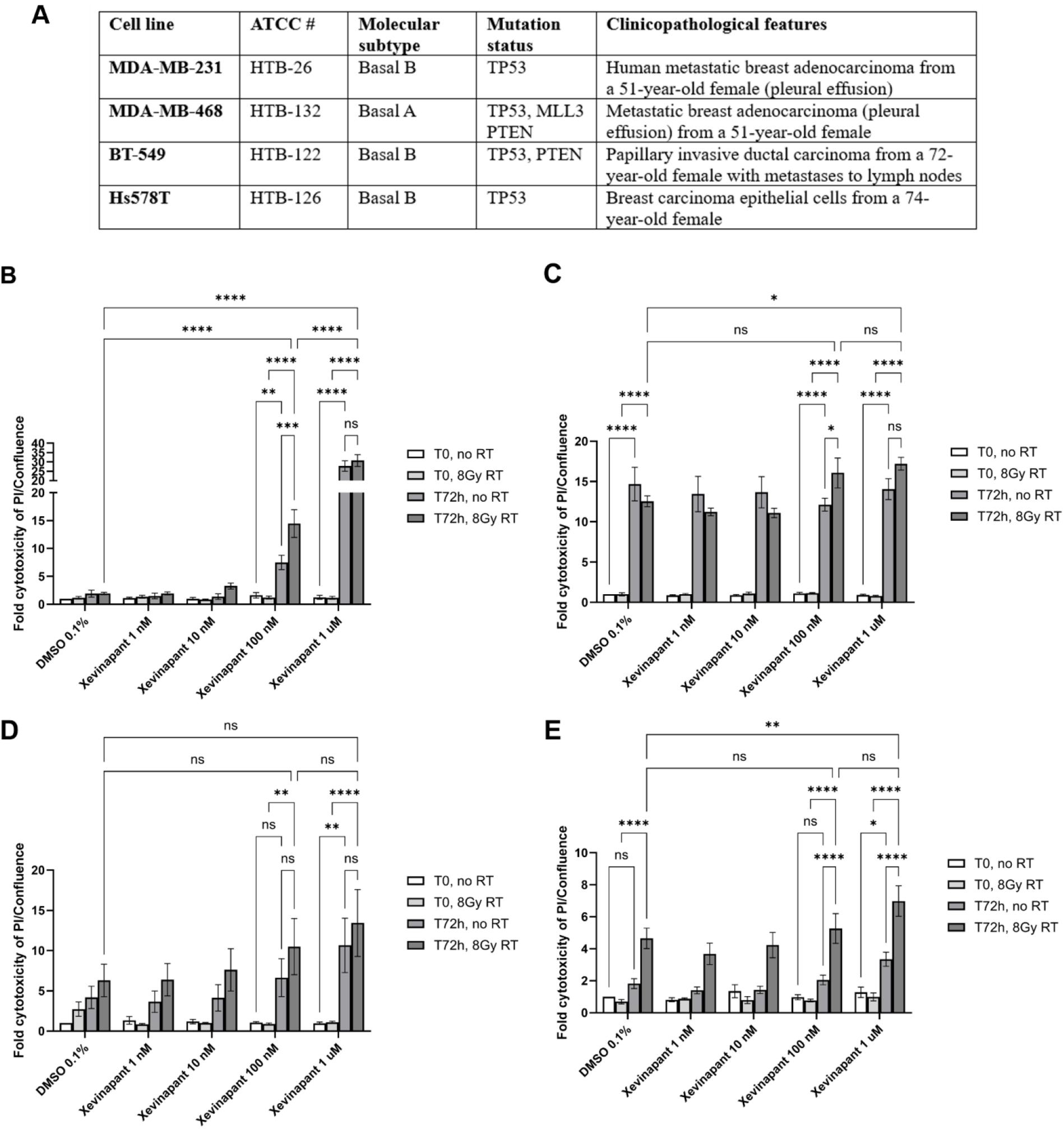
Xevinapant enhances radiation-induced cell death in a panel of TNBC cells. A. Panel of TNBC cells used as well as their molecular subtype, mutation status and clinicopathological features. B-E. MDA-MB231 (B), MDA-MB-468 (C), BT549 (D), Hs578T (E) cells were treated with increasing concentrations of Xevinapant (1nM to 1 μM) or control DSMO (0.1%) for 4 hours followed by RT at 8 Gy, and cell viability monitored over 72 hours by live cell imaging measuring propidium iodide incorporation into cells. Cell death was expressed as fold-cytotoxicity (ratio of PI-positive cells to % confluence area) normalized to control DMSO condition at T = 0. Results represent mean ± standard error (SE) of triplicates from 3 independent experiments (N = 9). Two-way ANOVA with Tuckey’s multiple comparison was performed, ns (not significant), ^*^ P ≤ 0.05, ^**^ P ≤ 0.01, ^***^ P ≤ 0.001, ^****^ P ≤ 0.0001

To get a better sense of the specific effect of Xevinapant on RT-induced cell death, we compared fold cytotoxicity at 72h in the presence and absence of RT. At a concentration of 100 nM, Xevinapant doubled MDA-MB-231 cytotoxicity in the presence of 8Gy RT compared to no RT (14-fold vs 7.5 fold at 72h, p = 0.0002) but not at 1 uM due to high sensitivity of the cells to Xevinapant alone at that concentration (30.8 fold vs 27.8 fold, ns). Likewise, Xevinapant increased RT-induced cytotoxicity in MDA-MB-468 at 100 nM (16-fold vs 12-fold, p = 0.0348) but not at 1 uM (17-fold vs 14-fold, ns), and in HS578T at both 100 nM (5-fold vs 2-fold, p <0.0001) and 1uM (7-fold vs 3-fold, p <0.0001). BT-549 did not show any significant increase in RT-induced cytotoxicity at 72h (10.4-fold vs 6.6-fold at 100 nM, and 13.4-fold vs 10.7-fold at 1uM, ns). Altogether these results support that Xevinapant enhances RT-induced cytotoxicity of TNBC cell lines in a concentration- and time-dependent manner. The magnitude of this effect varied across cell lines, with MDA-MB-231 being the most sensitive to combined Xevinapant and RT-induced cytotoxicity. MDA-MB-231 was therefore selected for subsequent experiments to further characterize the mechanisms underlying the interaction between Xevinapant and RT.

### Effect of Xevinapant on clonogenic survival following RT

Clonogenic survival assay is the gold standard for establishing radiosensitization.^21,22^ Thus, to confirm a radiosensitizing effect of Xevinapant in TNBC, we treated MDA-MB-231 cells with 1 uM of Xevinapant for 4 hours followed by irradiation at different doses and measured clonogenic survival after 14 days. We did not find any significant effect on clonogenic survival with a single dose of Xevinapant for the course of the experiment (Figure 2A, continuous condition). Matzinger *et al* previously showed that Xevinapant (also known as Debio 1143 or AT-406) only caused radiosensitization of HNSCC cells when administered continuously or added 24 h post-irradiation and maintained for at least 10 days.^23^ To simulate a more continuous supply of the drug *in vitro*, cells were pretreated with Xevinapant or control DMSO for four hours, then treated with RT and media replaced every three days with fresh Xevinapant at 1uM concentration (q3d condition). Xevinapant caused a modest but significant radiosensitization of MDA-MB-231 under these conditions, with a dose-enhancing factor (DEF) of 1.17 at a surviving fraction of 0.1 (Figure 2B). There was no radiosensitizing effect when cells were pre-treated for two hours followed by RT (Figure 2C). These results suggest that Xevinapant effect on the clonogenic survival of MDA-MB-231 cells is schedule-dependent and prolonged exposure to the drug is necessary to achieve a significant effect.

**Figure 2.**
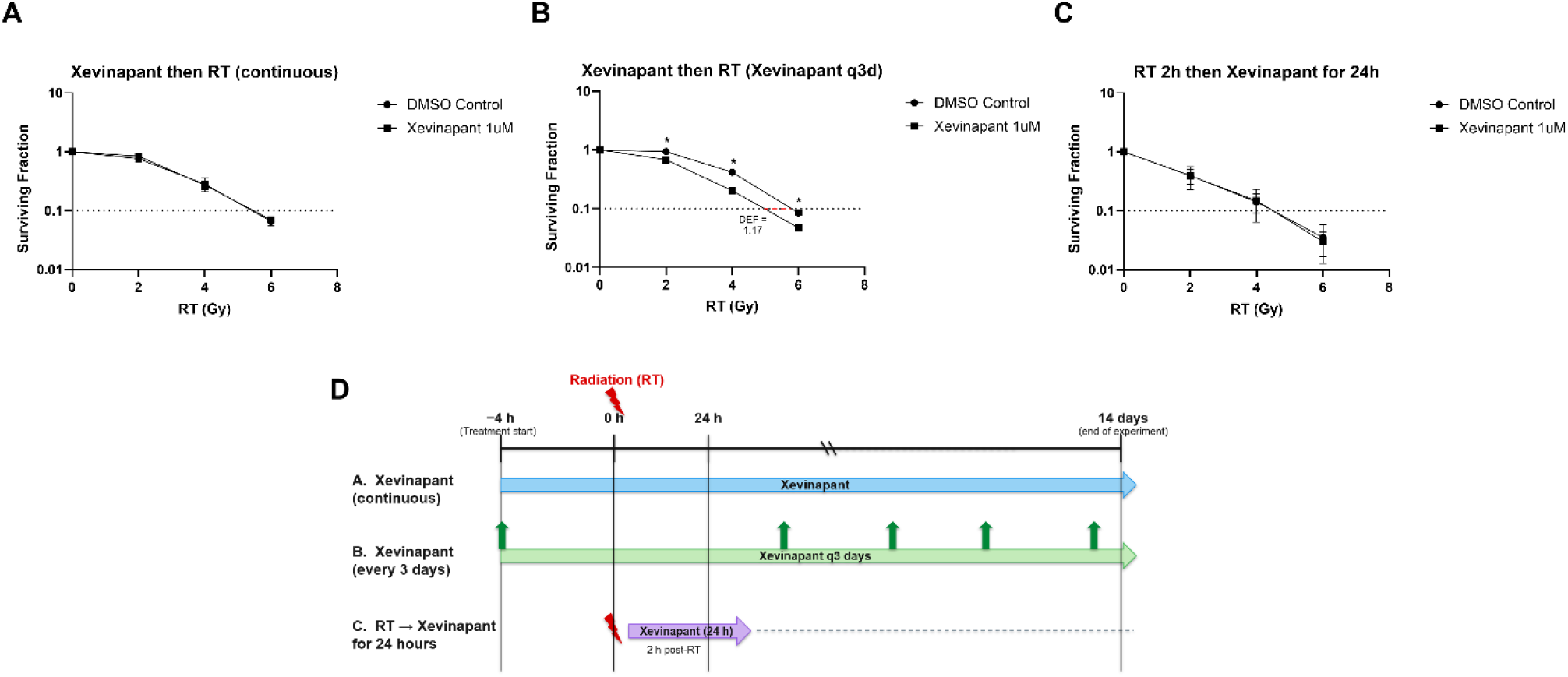
Effect of Xevinapant on clonogenic survival of TNBC cells following RT. A. MDA-MB-231 cells were treated with 1 μM Xevinapant or control DMSO (0.1%) for 4 hours then treated with different doses of RT (0, 2, 4, 6 Gy) and colony formation assessed after 14 days with no change in growth media (continuous condition). B. MDA-MB-231 cells were treated as above and media replaced every three days with fresh Xevinapant at 1uM concentration or control DMSO 0.1% (q3d condition). C. MDA-MB-231 cells were treated with RT first for 2 hours followed by treatment with Xevinapant 1 uM or DMSO 0.1% for 24 hours before the media was changed and colony formation assessed. D. Schematic showing the three different treatment schedules. In all three conditions, the dose enhancement factor (DEF) at a surviving fraction of 0.1 was calculated as the ratio of the RT dose required to achieve 10% survival in DMSO-treated cells to that required in Xevinapant-treated cells. A DEF >1 was considered indicative of radiosensitization by Xevinapant. Results represent mean ± standard error (SE) of triplicates from 3 independent experiments (N = 9). Statistical significance was assessed using multiple t-tests, with P values adjusted for multiple comparisons using the Holm–Šídák method, ^*^ P ≤ 0.05.

### Xevinapant radiosensitization of MDA-MB-231 is TNFα-dependent

To further characterize the potential radiosensitization properties of Xevinapant in the TNBC cell line MDA-MB-231, we set out to explore the role of TNFα and of apoptosis. Indeed, SMs such as Xevinapant are known to induce apoptosis through degradation or inactivation of the IAPs, in particular XIAP.^10,11^ Furthermore, TNFα plays an important role in SMs-induced apoptosis through binding to its receptors, depletion of cIAP1/2 and activation of the extrinsic apoptosis pathway.^8,9^ To investigate this, NLS-GFP stable cells expressing fluorescent GFP were established to accurately cell numbers by live cell imaging.

NLS-GFP/MDA-MB-231 were pretreated with 100 nM of Xevinapant alone or in the presence of exogenous human recombinant TNFα for four hours prior to being irradiated, and cytotoxicity measured over time through PI incorporation. Cytotoxicity was expressed as the ratio of PI positive cells to total number of cells expressing GFP. As shown earlier, Xevinapant increased RT-induced cytotoxicity and this was significantly enhanced by the addition of TNFα 10 ng/ml (6.9-fold vs 3.5-fold, p = 0.02) (Figure 3A). To assess specifically the role of TNFα in Xevinapant radiosensitization, clonogenic assays were repeated in native MDA-MB-231 in the presence of Xevinapant and TNFα for four hours followed by RT and colony formation assessed after 14 days. TNFα significantly enhanced Xevinapant-induced radiosensitization with a DEF of 1.57 (Figure 3B and C), compared to 1. 17 for Xevinapant only (Figure 2B). To assess whether the radiosensitization observed is apoptosis dependent, clonogenic assays were repeated in the presence of the pan-caspase inhibitor zVAD-fmk. TNFα or zVAD-fmk on their own did not decrease clonogenic survival, however TNFα-dependent radiosensitization was completely abrogated by the addition of zVAD-fmk (Figure 3D, P ≤ 0.0001 for Xevinapant + TNFα compared to Xevinapant + TNFα + zVAD). Taken together, these results suggest that Xevinapant-induced radiosensitiztion is indeed TNFα- and apoptosis-dependent.

**Figure 3.**
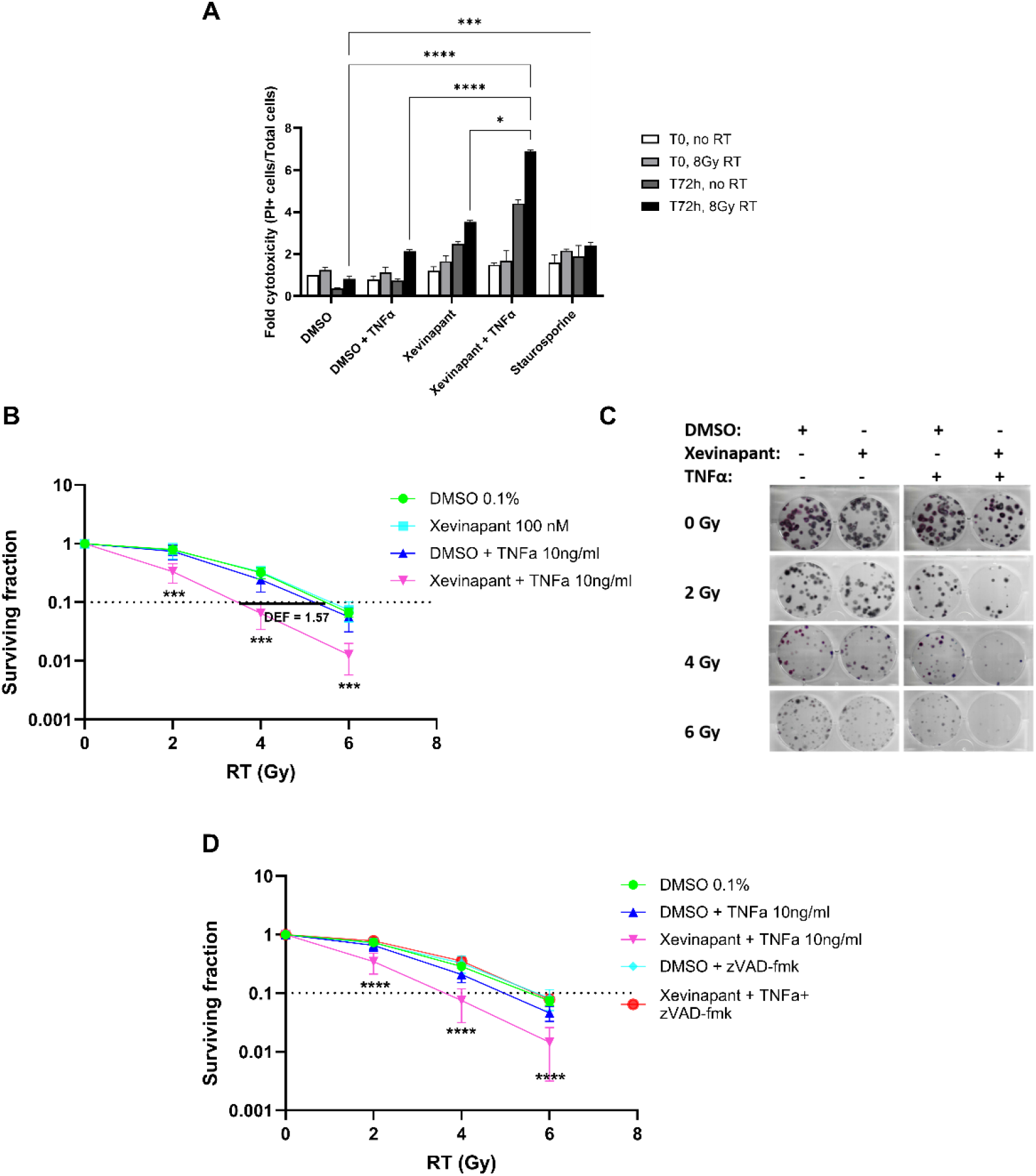
Xevinapant sensitizes MDA-MB-231 cells to radiotherapy in a TNFα-dependent manner. A. NLS-GFP/MDA-MB231 cells were treated with Xevinapant 100 nM or control DMSO +/-recombinant human TNFα 10 ng/ml for 4 hours before being treated with RT at 8 Gy. Cell cytotoxicity was expressed as the ratio of PI positive cells to total number of cells expressing GFP, normalized to control (DMSO at time 0). B. MDA-MB-231 were pretreated as above prior to being treated with different RT doses (0, 2, 4, 6 Gy) and colony formation assessed after 14 days. DEF was calculated as the ratio of the RT dose required to achieve 10% survival in DMSO-treated cells compared to Xevinapant+TNFα-treated cells. C. Representative number of colonies per treatment condition, showing significant decrease in colony formation in the Xevinapant+TNFα irradiated groups. D. MDA-MB-231 cells were treated as in B, with the addition of the pan-caspase inhibitor zVAD-fmk at 20 μM and colony formation assessed. Results represent mean ± standard error (SE) of triplicates from 3 independent experiments. Two-way ANOVA with Tuckey’s multiple comparison for the cell cytotoxicity assays and multiple t-tests adjusted for multiple comparisons using the Holm–Šídák method for clonogenic assays. ^*^ P ≤ 0.05, ^**^ P ≤ 0.01, ^***^ P ≤ 0.001, ^****^ P ≤ 0.0001.

### Exploring the mechanism of action of Xevinapant radiosensitization in TNBC cells

To further characterize the mechanism of Xevinapant-induced radiosensitization, MDA-MB-231 were treated with Xevinapant 100 nM and RT 4Gy in the presence or absence of TNFα 10 ng/ml and western blotting done to analyze protein markers of apoptosis and DNA damage at different timepoints (Figure 4A). Xevinapant notably decreased cIAP1 and XIAP protein levels, and to a lesser extent cIAP2. Cleaved PARP levels increased in combined Xevinapant + TNFα treatment at 24 hours compared to treatment with Xevinapant or TNFα alone. This differential increase in cleaved PARP levels persisted at 48 hours and 72 hours, consistent with delayed induction of apoptosis. γH2AX protein levels slightly increased at 24 hours in the combination group but increased in all condition groups in later time points, particularly at 72 h, suggesting persistent RT-induced DNA damage rather than a Xevinapant specific effect. Active caspase-3 staining was performed too but the protein levels were too faint to be reliably detected. As an alternative, we measured fluorescent activated cleaved caspase 3 and 7 levels in cells over time using live cell imaging. Xevinapant induced caspase-3/7 activation only in the presence of TNFα and RT (Figure 4B), confirming that TNFα is necessary for Xevinapant-induced apoptosis in irradiated MDA-MB-231.

**Figure 4.**
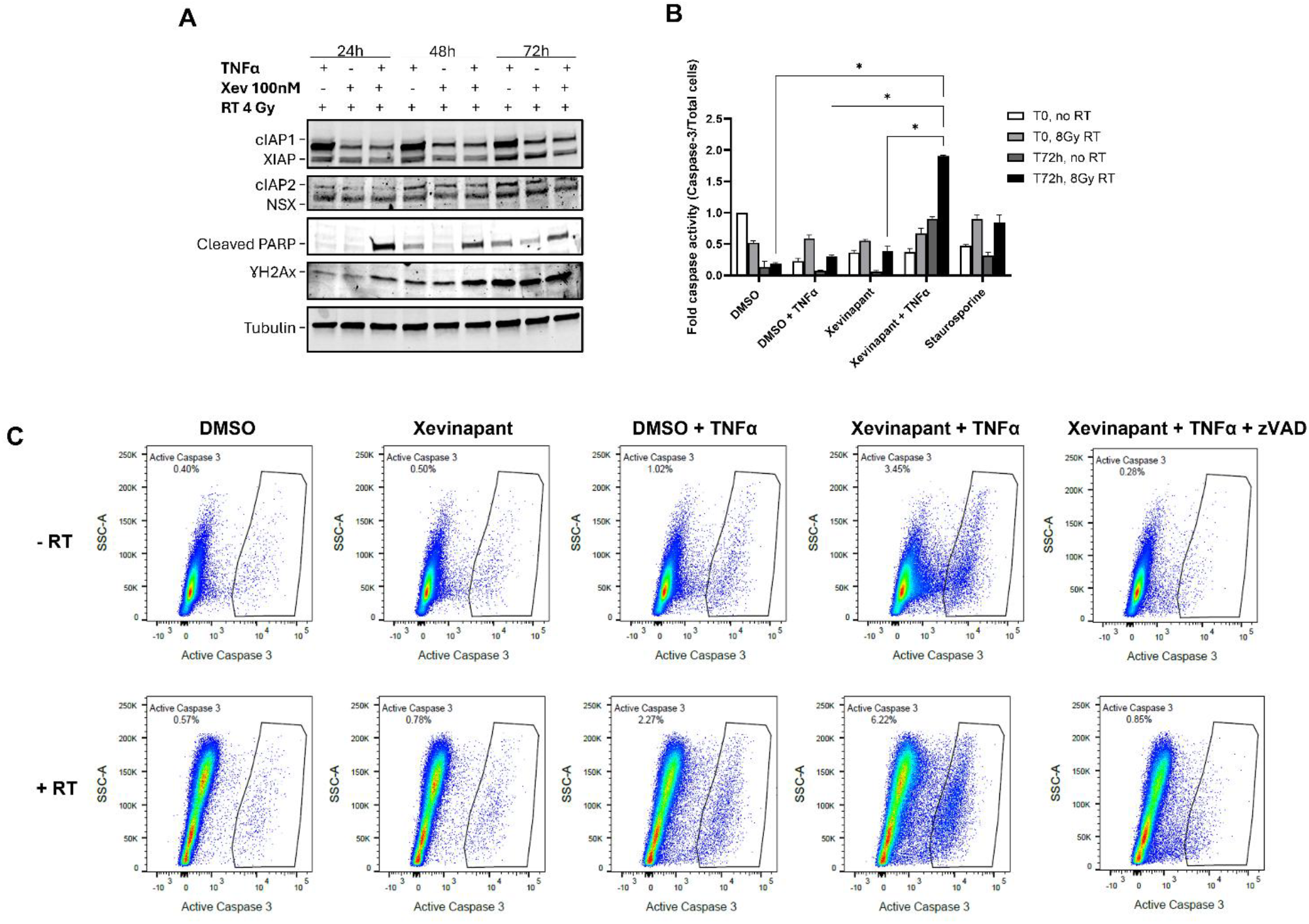
Xevinapant radiosensitization of MDA-MB-231 is TNFα- and apoptosis-dependent. A. MDA-MB-231 were treated with Xevinapant 100 nM (+) or control DMSO (-) +/-recombinant human TNFα 10 ng/ml for 4 hours before being treated with RT at 4 Gy. Cells lysates were collected at 24, 48 and 72 hours post RT and analyzed by western blot for protein expression of cIAP1, cIAP2, XIAP, phospho-cleaved PARP (Asp214) and phosphor-γH2Ax. Tubulin was used as a loading control, NSX = nonspecific band. B. NLS-GFP/MDA-MB-231 were treated as in A and RT 8 Gy, caspase-3/7 activity measured over time by live cell imaging and fold caspase activity expressed as the number of cells positive for activated Caspase-3/7 by the total number of cells expressing GFP, normalized to control (DMSO at time 0). C. Cells were treated as in A and flow cytometry analyses done at 48 hours to measure caspase activity using a specific BV650-conjugated active caspase-3 antibody.

To further characterize the apoptotic response at a single-cell level, we quantified the proportion of cells expressing intracellular active caspase-3 by flow cytometry following treatment with Xevinapant, TNFα and RT alone or in combination. RT, Xevinapant or TNFα alone did not increase the amount the cells entering apoptosis. However, the combination of RT, Xevinapant and TNFα increased the number of MDA-MB-231 cells entering apoptosis (6.22%) compared to the combination of Xevinapant and TNFα (3.45%) (Figure 4C). Though these numbers appear small, they represent the proportion of cells entering apoptosis at any given time. In another set of experiments where both ‘‘attached’’ and ‘‘floating’’ cells were collected and assessed by flow cytometry to account for the total number of cells that underwent apoptosis over 72 hours, the proportion of active caspase-3 expressing cells was 41.2% in the combined RT, Xevinapant and TNFα group compared to 16.9% with Xevinapant + TNFα, 20.3% with TNFα + RT, 9% with Xevinapant +RT and 5.8% with TNFα alone (Supplementary figure 1). In both cases, the proportional increase of apoptotic cells observed by flow cytometry is consistent with the fold-apoptosis increase observed in the live cell imaging experiments (Figure 4B). We next looked at the effect of the combination treatment on the cell cycle through PI incorporation into DNA and flow cytometry. RT treatment increased the proportion of cells in G2/M at four or 24 hours compared to non-irradiated cells but the addition of Xevinapant and TNFα did not increase this (Supplementary Figure 2). Altogether, these results show that Xevinapant sensitizes MDA-MB-231 to RT-induced apoptosis in a TNFα-dependent manner.

## Discussion

There is an unmet need in the treatment of TNBC as recurrence rates and progression to metastatic disease can be as high as 50% in patients that did not achieve a pathological complete response (pCR),^4^ thus novel treatment strategies are warranted to improve outcomes for patients with TNBC. In this study, we investigated whether pharmacologic antagonism of the inhibitor of apoptosis proteins (IAPs) by Smac mimetics (SMs) could enhance TNBC cells’ response to radiotherapy (RT). We show that the SM Xevinapant enhances radiation-induced cell death across multiple TNBC cell lines, although the magnitude of this effect varies between cell lines. In MDA-MB-231 cells, which showed the strongest response to combined treatment, Xevinapant-mediated radiosensitization was dependent on treatment schedule and was markedly potentiated by TNFα. Mechanistically, the combination was associated with depletion of IAP proteins, increased PARP cleavage and caspase-3/7 activation, while inhibition of caspase activity completely abrogated the clonogenic radiosensitizing effect. Collectively, these findings support a model in which Xevinapant lowers the apoptotic threshold of irradiated TNBC cells, with TNFα acting as an important mediator of the response.

Our findings are corroborated by previous observations that SMs can enhance radiation-induced cell death as they promote cIAP1/2 degradation and antagonize XIAP-mediated caspase inhibition, thereby facilitating apoptosis in response to cellular stress.^6,10,11^ Indeed, previous studies using the bivalent Smac mimetic SM-164 demonstrated caspase-dependent radiosensitization in both head and neck and breast cancer cells, with sensitization dependent on caspase activation.^16,17^ Our results with Xevinapant in TNBC are consistent with this mechanism and identify TNFα as a major modulator of the interaction between Xevinapant and radiation.

Interestingly, Xevinapant’s radiosensitizing effect depended strongly on the treatment schedule (Figure 2). Continuous exposure without drug replenishment produced little effect on long-term clonogenic survival, whereas replenishment of Xevinapant every three days resulted in radiosensitization. Conversely, administering Xevinapant for only 24 hours after irradiation was insufficient to substantially alter clonogenic survival. This schedule dependence is consistent with previous work in head and neck cancer demonstrating that prolonged Xevinapant exposure was required for optimal radiosensitization.^23^

The most striking radiosensitivity enhancement occurred in the presence of TNFα. Addition of exogenous TNFα increased the DEF0.1 from 1.17 with Xevinapant alone to 1.57 with Xevinapant and TNFα, while TNFα alone did not substantially alter clonogenic survival (Figure 3). Importantly, with the presence of TNFα, only 100 nM of the drug was necessary to have a significant radiosensitizing effect. This finding is consistent with the established biology of SMs. cIAP1 and cIAP2 are critical regulators of TNF receptor signaling, and their depletion following SM treatment can redirect TNFα signaling toward formation of a death-inducing signaling complex and caspase-dependent apoptosis.^8,10,11^ Previous studies from our group and others have similarly demonstrated that the antitumor activity of SMs can depend on autocrine or microenvironmental TNFα.^12-14^ Matzinger *et al*. specifically also demonstrated that Xevinapant-mediated radiosensitization in head and neck cancer was TNFα-dependent,^23^ suggesting that the interaction identified here in TNBC may represent a conserved mechanism across tumor types rather than a breast cancer-specific phenomenon.

Multiple complementary experiments in our study support apoptosis as a major downstream mechanism of this interaction (Figure 3, 4, Suppl. Figure 1). The combination of Xevinapant, TNFα, and radiation was associated with increased PARP cleavage and caspase-3/7 activity, while flow cytometry showed a higher proportion of active caspase-3-positive cells. When both adherent and detached cells were recovered, active caspase-3 positivity reached 41.2% after the triple combination, compared with 16.9% after Xevinapant plus TNFα and 9% after Xevinapant plus RT. Most importantly, the pan-caspase inhibitor zVAD-fmk abrogated TNFα-enhanced radiosensitization, demonstrating the functional importance of caspase activation. This functional rescue provides stronger evidence for a causal role of caspase-dependent cell death and is consistent with previous studies in breast and head and neck cancer in which inhibition of caspases reduced SM-mediated radiosensitization.16,17 ^16,17^ In contrast, γH2AX protein was not selectively increased by Xevinapant at later time points, suggesting that Xevinapant and TNFα radiosensitization was not primarily through increased DNA damage. Furthermore, RT induced the expected accumulation of cells in G2/M, whereas addition of Xevinapant and TNFα did not further increase this effect. These observations favor a model in which Xevinapant modifies the cellular response to radiation damage rather than substantially increasing the initial burden of DNA lesions or altering radiation-induced cell-cycle arrest.

Importantly, while preclinical and clinical data is limited in breast cancer, two phase I trials have demonstrated the safety and tolerability of the SM LCL161 as a single agent or in combination with Paclitaxel in breast cancer.^24,25^ Additionally, In a randomized phase II trial of 209 patients with TNBC, Bardia *et al*.^26^ tested the combination of neoadjuvant LCL161 with paclitaxel versus paclitaxel alone.

The primary endpoint was efficacy of neoadjuvant LCL161 in combination with paclitaxel, as defined by the pathologic complete response (pCR) rate at 12 weeks. Interestingly, patients with a TNFα-based gene signature (GS-positive patients) presented a significantly higher pCR rate with LCL161 and Paclitaxel compared to control (38.2% vs 17.2%) whereas GS-negative patients experienced lower pCR rates (5.6% vs 16.4%).^26^ There was however a higher incidence of grade 3 or 4 adverse events in the combination arm compared to the control arm, notably neutropenia (24.5%) and diarrhea (5.7%).^26^ Although this study evaluated chemotherapy rather than radiotherapy, the dependence of clinical response on a TNFα-related tumor phenotype parallels our preclinical findings and raises the hypothesis that TNFα pathway activity could serve as a predictive biomarker for SM-based radiosensitization. Furthermore, clinical experience with Xevinapant itself emphasizes the importance of patient selection and treatment context. In a randomized phase II trial investigating the SM Xevinapant plus chemoradiotherapy (CRT) vs. placebo plus CRT in 96 patients with unresected locally advanced head and neck cancer squamous cell cancer (HNSCC), the risk of death was decreased by half in the Xevinapant arm compared with placebo (adjusted HR 0.47; 95% CI, 0.27–0.84; P = .0101) while incidence of late-onset grade ≥3 toxicities was similar across arms^18^. However, these findings were not confirmed in the subsequent phase III TrilynX trial in which Xevinapant plus chemoradiotherapy did not improve event-free survival in 730 randomized patients. Event-free survival was shorter in the Xevinapant arm, overall survival was unfavorable, and grade ≥3 and serious treatment-emergent adverse events were more frequent with Xevinapant.^27^ These findings further emphasize the need to better understand the type of patients that may response to combination therapy with Xevinapant and RT.

Several limitations of the present study should be acknowledged. The experiments were performed *in vitro* and predominantly performed in MDA-MB-231 cells. Exogenous TNFα was used to establish mechanistic dependence but does not reproduce the spatial and temporal complexity of TNFα production within the tumor microenvironment. Thus, the dependence on TNFα and caspase activation should be validated with *in vivo* syngeneic models of TNBC. Finally, although pharmacologic caspase inhibition strongly supports a role for apoptosis, additional genetic approaches targeting specific components of the TNFα-IAP-caspase pathway would further define the molecular determinants of response.

## Conclusions

Our findings demonstrate that Xevinapant can enhance radiation-induced cell death in TNBC and identify TNFα-dependent, caspase-mediated apoptosis as a central determinant of this interaction. These results are promising for the combined use of RT and Xevinapant to increase treatment outcomes in TNBC, and investigations are underway to further characterize the mechanism of Xevinapant radiosensitization in vivo and define a TNFα-related gene signature to identify patients with TNBC that could potentially respond to Xevinapant and radiotherapy.

## Supplementary Materials

Suppl. Figure 1, Suppl. Figure 2

## Author Contributions

Conceptualization, M.D.F.; methodology, M.D.F. and J.L.; software, M.D.F. and J.L.; formal analysis and validation, M.D.F.; resources, M.D.F., C.M. and P.W.; writing - original draft preparation, M.D.F.; writing - review and editing, M.D.F., J.L., P.W. and C.M.; supervision, P.W. and C.M. All authors have read and agreed to the published version of the manuscript.

