## Supplementary data for "The Smac mimetic drug Xevinapant sensitizes triple negative breast cancer to radiotherapy-induced cell death in a TNFα dependent manner"

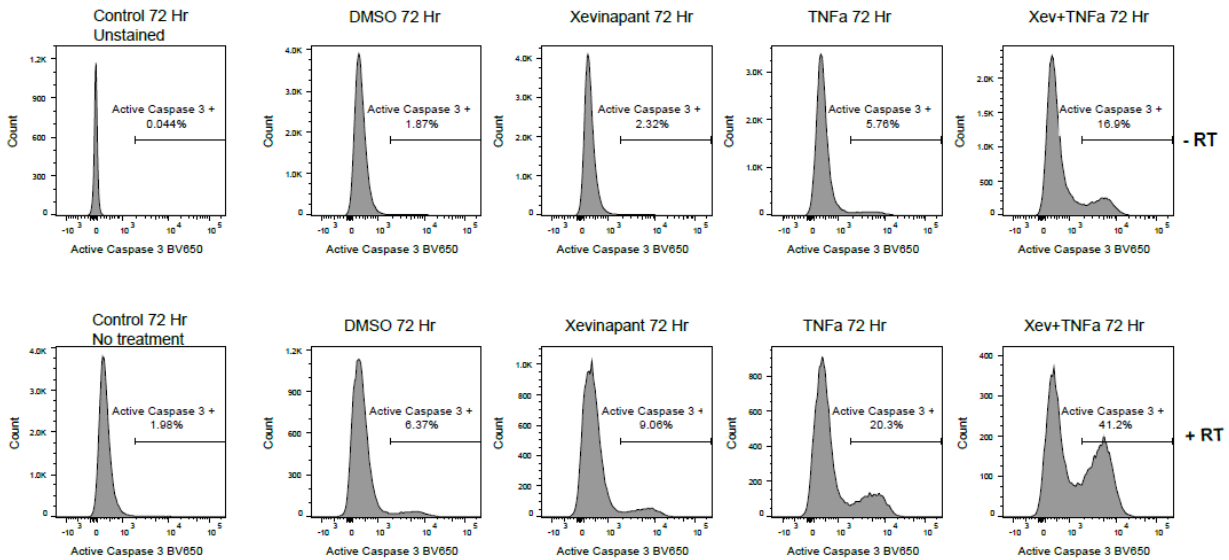

Supplementary Figure 1: Flow cytometry analysis of Caspase 3 activity in MDA-MB-231 treated with Xevinapant, TNFα and RT

MDA-MB-231 were treated with Xevinapant 100 nM (+) or control DMSO (-) +/- recombinant human TNFα 10 ng/ml for 4 hours before being treated with RT at 4 Gy and flow cytometry analyses done at 72 hours to measure caspase activity using a specific BV650-conjugated active caspase-3 antibody. Both floating and adherent cells were included in the analysis to capture the total amount of cells that underwent apoptosis during the course of the experiment.

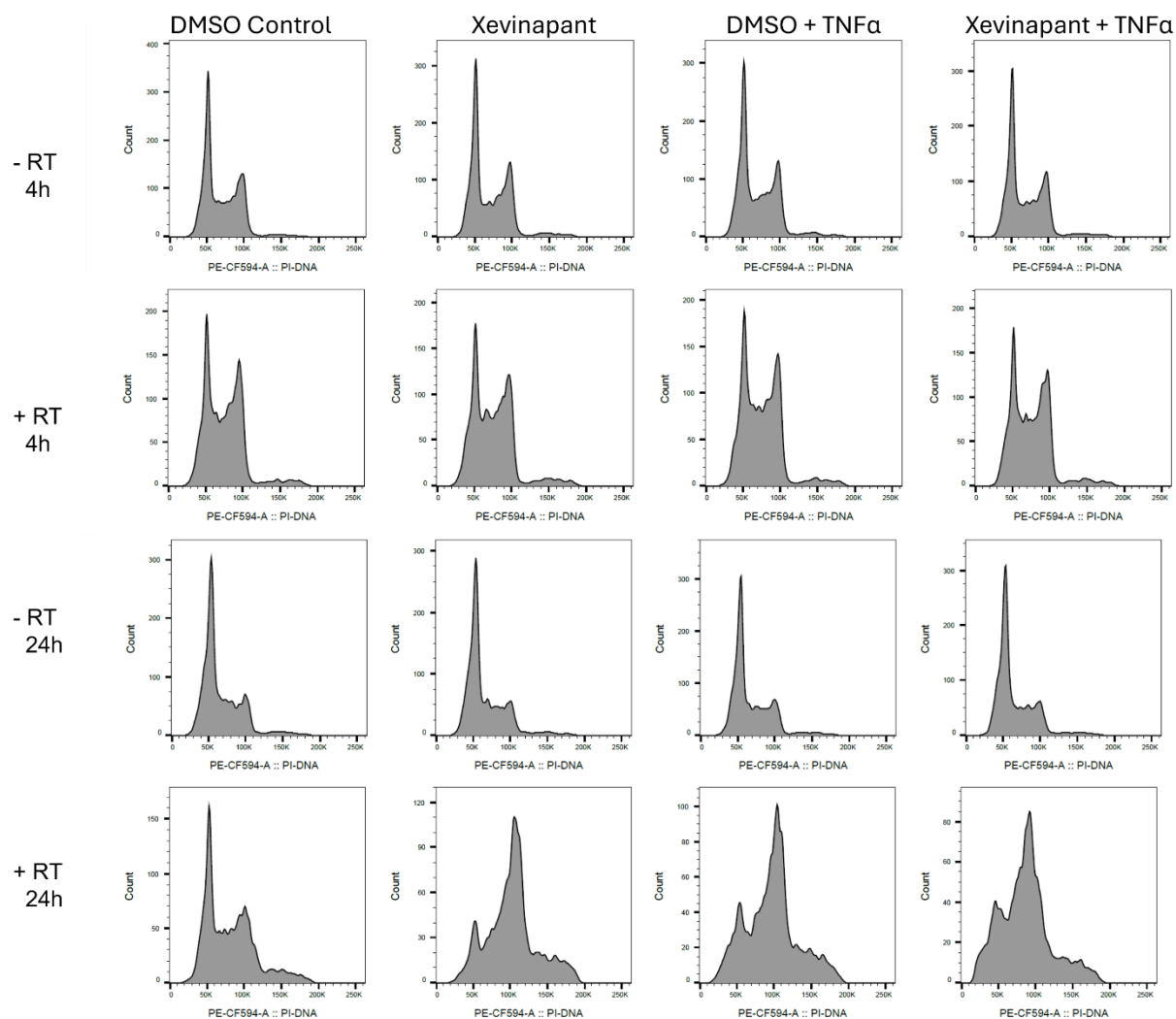

Supplementary Figure 2: Flow cytometry analysis of the cell cycle in MDA-MB-231 treated with Xevinapant, TNFα and RT

MDA-MB-231 were treated with Xevinapant 100 nM (+) or control DMSO (-) +/- recombinant human TNFα 10 ng/ml for 4 hours before being treated with RT at 4 Gy and flow cytometry analyses done at 4h and 24h to determine cell cycle distribution by propidium iodide incorporation
